# Elicitors of plant defense induce the accumulation of amino acids that suppress both bacterial virulence and growth

**DOI:** 10.1101/2022.05.30.494047

**Authors:** Xiaomu Zhang, Phil Tubergen, Israel Agorsor, Pramod Khadka, Connor Tempe, Eva Collakova, Guillaume Pilot, Cristian H. Danna

**Affiliations:** Department of Biology, University of Virginia, Charlottesville, VA 22904, USA; School of Plant and Environmental Sciences, Virginia Tech, Blacksburg, VA 24060, USA

## Abstract

Plant immunity relies on the perception of Microbe-Associate Molecular Patterns (MAMPs) from invading microbes to induce defense responses that suppress attempted infections. It has been proposed that MAMP-triggered immunity (MTI) suppresses bacterial infections by suppressing the onset of bacterial virulence. However, the mechanisms by which plants exert this action are poorly understood. Here we uncover that MAMP-perception in Arabidopsis induces the accumulation of free amino acids (AA) in a salicylic acid (SA) dependent manner. When co-infiltrated with Glutamine and Serine, two of the MAMP-induced highly accumulating amino acids, *Pseudomonas syringae* pv. tomato DC3000 expressed low levels of virulence genes and failed to produce robust infections in otherwise susceptible plants. When applied exogenously, Glutamine and Serine suppressed bacterial virulence and bacterial growth directly without the need for MAMP-perception and SA-signaling, bypassing MAMP-elicited defense. In addition, an increased level of endogenous Glutamine in the leaf apoplast of a gain-of-function mutant of *Glutamine Dumper-1* rescued the partially compromised bacterial virulence and growth suppressing phenotype of the SA-induced deficient-2 (*sid2*) mutant. Our data suggest that MTI suppresses bacterial infections by the direct suppressing effect of accumulating amino acids on the onset of bacterial virulence.

## INTRODUCTION

Plants are remarkably resistant to microbial infections. Only specialized microbes with evolved mechanisms to suppress inducible plant defense responses can cause disease (Abramovitch and Martin, 2004). Leaf bacterial pathogens initiate infections by breaching the stomatal barrier at the leaf surface and gaining access to the leaf apoplast, the leaf mesophyll’s water and nutrient-rich intercellular space (Xin and He, 2013). In the leaf apoplast, invading bacteria spend the first five hours without growing and adapting their metabolism to the new and rapidly changing apoplastic environment (Xin and He, 2013). Early in the interaction, both plant and bacterial cells perceive each other and activate transcriptional and metabolic programs tailored to gain control over each other (Preston, 2017). While plant responses evolved to suppress infection attempts, bacterial responses evolved to allow rapid adjustment to the leaf apoplastic environment, suppress plant defense responses, and promote leaf colonization (Yu et al., 2013; O’Leary et al., 2016).

Early in the interaction, plants recognize invading bacteria by their unique suite of molecules or activity patterns, generally known as Microbe-Associated Molecular Patterns (MAMPs). Plant-encoded Pattern-Recognition Receptors (PRR) perceive MAMPs at the cell surface and activate defense responses collectively known as MAMP-Triggered Immunity (MTI; also known as Pattern-Triggered Immunity or PTI) (Gómez-Gómez et al., 1999; Nicaise et al., 2009). At the cellular level, MTI early responses include the activation of Mitogen-Activated Protein-Kinases, cytoplasmic Ca^+2^ influx, production of reactive oxygen species, and genome-wide transcriptional reprogramming (Bigeard et al., 2015). Late MTI responses include the synthesis and accumulation of defense signaling molecule salicylic acid (SA) and ethylene (Gómez-Gómez et al., 1999; Asai et al., 2002; Lecourieux et al., 2005), the synthesis and accumulation of amino acid-derived secondary metabolites, the closing of the stomata, and the reinforcement of plant cell walls via callose and lignin deposition (Adams-Phillips et al., 2009; Clay et al., 2009; Chezem et al., 2017). Ultimately, MTI responses suppress bacterial growth via several mechanisms that act synergistically and coordinately. Suppressing MTI is a critical step in colonizing the plant host (Abramovitch and Martin, 2004). Non-pathogenic bacteria fail to suppress MTI responses, cannot adjust their metabolism to the rapidly changing leaf apoplast environment, and succumb within the first hours of the attempted colonization (Hauck et al., 2003; Clay et al., 2009). Bacterial pathogens, however, have evolved virulence mechanisms to suppress MTI early in the interaction (Xin et al., 2018). After entering the leaf apoplast, *Pseudomonas syringae* pv. tomato DC3000 (*Pst*DC3000) perceives Arabidopsis metabolites and induces the synthesis of Type-3 Secretion System (T3SS) secretory proteins, secreted T3SS effector proteins (T3E), and the SA-antagonist molecule coronatine (COR), all of which have a concerted activity suppressing plant defense (Bender et al., 1987; Brooks et al., 2005; Zhang et al., 2008; Xin and He, 2013).

Recent studies have shown that Arabidopsis plants that had been exposed to MAMPs and allowed to proceed with the MTI program for 24 h, suppressed the induction of *Pst*DC3000 T3SS, T3E, and COR biosynthesis genes (Lovelace et al., 2018; Nobori et al., 2018; Smith et al., 2018), suggesting that MTI suppresses bacterial growth via suppressing virulence. The MTI-mediated suppression of bacterial virulence seems to rely on changes in the metabolite composition of the leaf apoplast. Arabidopsis seedlings exposed to elf26, a 26aa long synthetic peptide containing the minimal epitope of bacterial translation Elongation Factor-Tu, suppressed the onset of *Pst*DC3000 virulence via lowering the availability of citric acid, aspartic acid, and 4-benzoic acid (Anderson et al., 2014). Similarly, the perception of flg22, a 22aa long synthetic peptide containing the minimal epitope of bacterial flagellin, induced changes in sugar concentrations in the leaf apoplast of Arabidopsis that compromised the onset of *Pst*DC3000 virulence (Yamada et al., 2016). Overall, these studies suggest that MAMP perception elicits changes in the concentration of plant metabolites in the leaf apoplast that affect bacterial virulence and growth.

In addition to sugar polymers, secreted proteins, and complex secondary metabolites, primary plant metabolites such as hexoses and free amino acids (hereinafter referred to as “AA”), are among the most abundant metabolites in the Arabidopsis leaf apoplast. Notably, some of these metabolites (e.g., glucose and several AA) suffice to support *Pst*DC3000 growth and regulate the expression of *Pst*DC3000 virulence genes *in vitro*, suggesting a critical function for these metabolites in controlling bacterial infections (Rahme et al., 1992; Rico and Preston, 2008; Park et al., 2010; Anderson et al., 2014; Chatnaparat et al., 2015a; Chatnaparat et al., 2015b; Yamada et al., 2016). In addition, organic acids, gamma-aminobutyric acid (GABA), and some AA accumulate at lower millimolar concentrations in *Cladosporium fulvum*-infected tomato leaf (Solomon and Oliver, 2001; Solomon and Oliver, 2002). Similarly, GABA and several proteogenic AA accumulate to high concentrations in the leaf of *P. syringae*-infected Arabidopsis and bean plants (Ward et al., 2010; O’Leary et al., 2016), suggesting that these metabolites could play a role in defense or susceptibility to pathogens.

In a previous study, we have presented evidence that MAMP-perception elicits changes in both, AA transport activity and extracellular AA concentrations. Importantly, high concentrations of extracellular AA boosted bacterial growth in mock-treated seedlings but suppressed bacterial growth when MTI was elicited 24 h prior to the inoculation (Zhang et al., 2022). The present study aimed to address the mechanisms by which flg22-elicited changes in AA concentrations impact the growth of *Pst*DC3000. Here we present evidence of a net accumulation of Gln and Ser, among other less represented AA, in MAMP-treated leaves and leaf apoplast of Arabidopsis plants. The MTI-induced increased concentration of AA depends on an intact SA-mediated signaling pathway and suffices to suppress *Pst*DC3000 virulence and growth, suggesting that MAMP-elicited accumulation of AA plays a pivotal role in suppressing bacterial infections. Overall, our data provide new insights into how MAMP perception leads to metabolite changes that confer enhanced immunity to bacterial infections.

## Results

### MAMP perception elicits the accumulation of amino acids in flg22-treated leaves

AA transport activity and the extracellular concentrations of AA dynamically change in response to MAMPs perception liquid-grown 10-days old Arabidopsis seedlings (hereafter referred as “seedlings”). Extracellular AA concentrations increase within 2-4 h post-treatment (HTP) and then drop between 8-24 HPT with MAMPs (Zhang et al., 2022). In agreement with our previously published data, the flg22-induced enhanced AA uptake activity led to a significant increase in AA concentration in flg22-treated compared to mock-treated seedlings (**Figure 1A**). To determine if, like in seedlings, the elicitation of MTI leads to changes in AA concentrations in treated leaves and leaf apoplast of six-week-old plants (hereafter referred to as “plants”), the concentration of AA was assessed in fully expanded mature leaves at 4, 8, 20, and 24 HPT with flg22. Like in whole seedlings, the concentration of AA in leaves started to increase by 20 HPT (**Figure 1B)**. In the leaf apoplast, the concentration of AA increased significantly in flg22-treated leaves 24 HPT (**Figure 1C**). The flg22-receptor mutant *fls2* did not accumulate AA in leaf tissue in response to flg22 treatment (**Figure S1A**), demonstrating that AA accumulation in treated leaves is a signaling-dependent process.

**Figure 1:**
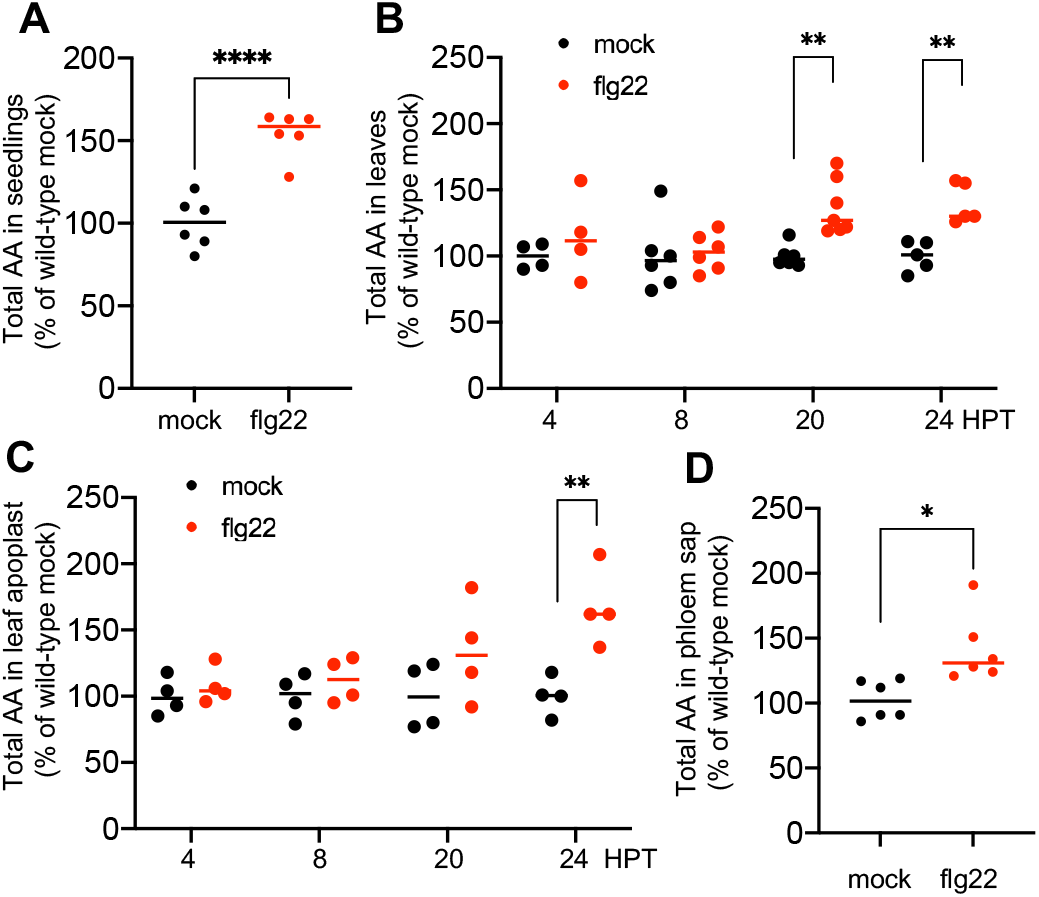
MAMP perception induces the accumulation of amino acids in treated leaves. **(A)** Concentrations of AA in eleven-day-old seedlings 24 h post-treatment with water (black) or flg22 (red). (**B**) Total AA in leaves of wild-type plants after infiltration of leaves with water (black) or flg22 (red). (**C**) Total AA in leaf apoplastic washing fluids of wild-type plants treated with water (black) or flg22 (red) 24 h post-treatment. (**D**) Total AA in the phloem sap of wild-type leaves 24 HPT. Statistic analysis: Welch t-test (A) and t-tests for (B)(C)(D)(E). All figures show one representative experiment from at least 3 independent experiments.

To define which individual amino acids contribute to increasing the concentrations of total AA following flg22 perception, leaves were mock-treated or infiltrated with flg22 and then collected for AA extraction and profiling 24 HPT. Among the individual AA detected in leaf samples, Ala, Asn, Gln, Glu, Gly, His, Phe, Ser, Val increased significantly compared to mock-treated samples (**Table I**). We found that the concentration of AA in phloem sap also significantly increased 24 HPT with flg22 (**Figure 1D**), suggesting that the increased concentration of total AA in treated leaves does not result from an inhibition of phloem loading. Hence, we hypothesized that increased activity of cellular processes that generate AA, or a decreased activity of processes that consume AA, could contribute to AA buildup in flg22-treated leaves. Based on the GO terms associated with amino acid metabolism (MapMan), we analyzed publicly available gene expression datasets obtained from flg22-treated vs mock-treated leaves. This analysis revealed that genes that contribute AA to the cellular pool (i.e., AA biosynthesis, autophagy, senescence, and proteolysis) are significantly induced 1 HPT and 4 HPT with flg22 (**Figure S2, Table S1**). Conversely, with the only exception of glucosinolates biosynthesis genes that were induced at 1 HPT and 4 HPT, genes involved in processes that consume AA (i.e., protein biosynthesis, amino acids catabolism) were upregulated at 1 HPT but downregulated at 4 HPT with flg22 (**Figure S2, Table S1**). These data suggest that, while metabolic changes that contribute AA to the cellular pool are induced, cellular processes that consume AA are downregulated by flg22 perception.

**Table 1.** Amino acids in leaves of wild-type plants 24 HPT

|  | mock |  | flg22 |  | Adjusted P Value |
| --- | --- | --- | --- | --- | --- |
|  | Mean | SEM | Mean | SEM |  |
| Ala | 142.9 | 7.3 | 216.3 | 5 | 2.44E-08 * |
| Arg | 6.8 | 0.4 | 7.3 | 0.4 | 8.49E-01 |
| Asn | 47 | 2.7 | 58.3 | 2.7 | 4.75E-02 * |
| Asp | 285 | 8.9 | 291.5 | 8.5 | 9.38E-01 |
| Gln | 277.9 | 16.8 | 352.7 | 19.1 | 4.75E-02 * |
| Glu | 342.3 | 14.2 | 527.9 | 15.8 | 6.10E-09 * |
| Gly | 38.2 | 2.8 | 96.8 | 8.3 | 1.07E-06 * |
| His | 4.7 | 0.2 | 8 | 0.6 | 9.48E-05 * |
| Ileu | 3.9 | 0.2 | 6.3 | 1 | 1.70E-01 |
| Leu | 5.9 | 0.6 | 9.9 | 1.8 | 1.97E-01 |
| Lys | 6.8 | 0.6 | 8.8 | 0.9 | 3.12E-01 |
| Phe | 5.5 | 0.3 | 12.7 | 1.1 | 3.44E-06 * |
| Pro | 29.6 | 3.9 | 28.2 | 4.9 | 9.38E-01 |
| Ser | 158 | 7.6 | 226.3 | 8.2 | 8.50E-06 * |
| Thr | 86.5 | 3.4 | 84.7 | 2.7 | 9.38E-01 |
| Val | 11.8 | 0.4 | 18.6 | 1.8 | 5.46E-03 * |

(\*) Statistically significant differences at $p \leq 0.05$ between wild-type mock- and flg22-treated leaves. Concentrations of free AA (pmol/mg DW) in extracts of water- (mock) or flg22-treated wild-type leaves 24 HAE. Mean ± SEM of 18 mock samples and 17 flg22 samples from 4 independent experiments. Data were analyzed with multiple Welch t-test; p values were adjusted by two-stage step-up false discovery rate method (Q = 0.01).

### Amino acids that accumulate in MTI-elicited leaves directly suppress bacterial growth and virulence bypassing plant responses

To elucidate the potential biological function underling the MAMP-induced increased AA concentration in plant immunity, we sought to test if and how individual AA modify *Pst*DC3000 growth in the leaf. A pilot experiment using radiolabeled AA showed that Pro, Gln, and Ser were quickly absorbed by leaf cells after infiltration of the amino acids in the apoplast (**Figure S3)**. This suggested that if amino acids were infiltrated in the leaf prior to bacterial infection, they would be rapidly taken up by the plant cells and made unavailable to *Pst*DC3000. Hence, AA had to be co-infiltrated with bacteria to ensure their availability, at least for a short period of time. To mimic the high availability of AA present in the apoplast of MTI-elicited plants, we initially tested the effect of two different concentrations of a mix of two of the flg22-induced accumulating AA, Gln+Ser, on *Pst*DC3000 growth. The data showed that the co-infiltration of *Pst*DC3000 with 10 mM Gln+Ser reproducibly inhibited bacterial growth compared to *Pst*DC3000 inoculated alone (**Figure S4**), while the results varied between inhibition or no effect for the 1 mM concentration, depending on the experiments. We thus decided to use 10 mM AA in future experiments. Interestingly, while co-infiltration with Gln, Ser or Val suppressed *Pst*DC3000 growth assessed 48 HPI, His boosted *Pst*DC3000 growth compared to non-supplemented bacteria (**Figure 2A**). Externally supplied Glu and Gln have been reported to induce SA signaling-mediated defense responses in Arabidopsis five days after treatment (Goto et al., 2020). If Gln+Ser supplementation had a plant-defense inducing activity, it could indirectly suppress bacterial growth by eliciting plant defense. To test this possibility, we assessed the expression of MTI and SA defense-related marker genes in plants that have never been treated to elicit defense responses (hereafter referred to as naïve plants). We infiltrated mature leaves of naïve plants with Gln+Ser or co-infiltrated *Pst*DC3000 with Gln+Ser to assess plant responses. Infiltration of leaves with Gln+Ser alone did not significantly induce the MTI marker genes *CYP81F2* and *WRKY29* (**Figure S5A**). However, the SA biosynthesis gene *ICS1* (*Iso-Chorismate Synthase-1*) and the SA defense marker gene *PR1* (*Pathogenesis Related-1*), were modestly but significantly induced 24 h after Gln+Ser infiltration (**Figure S5B**). These data suggest that Gln and/or Ser could prime plants for defense responses. In addition, *CYP81F2* showed modest but significantly higher expression in leaves infiltrated with *Pst*DC3000 and Gln+Ser compared to those infiltrated with *Pst*DC3000 alone (**Figure S5C**). Nevertheless, when Gln+Ser were co-infiltrated with *Pst*DC3000 into leaves of naïve wild-type plants, *ICS1* and *PR1* were expressed at the same level as those detected in leaves infiltrated with bacteria alone (**Figure S5D**). *CYP81F2* induced expression in leaves co-infiltrated with *Pst*DC3000 and Gln+Ser is likely due to the virulence suppressing effect of Gln+Ser on *Pst*DC3000. MAMP perception strongly induces *CYP81F2*, a gene needed for MAMP-induced callose deposition. However, *Pst*DC3000 suppresses *CYP81F2* expression and callose synthesis early in the infection (Clay et al., 2009). These data suggest that Gln+Ser supplementation could impair the ability of *Pst*D3000 to suppress early MAMP-elicited responses that would otherwise go undetected. To test this hypothesis, we examined *hrpL* gene expression, a major T3SS regulator. Following the conditions established in an early study (Rahme et al., 1992), we grew *Pst*DC3000 in *hrpL*-inducing minimal medium (HMM) supplemented with 5 mM of each AA for 150 min before collecting samples. Except for Phe, every AA tested decreased *hrpL* expression *in vitro*, with Gln and Ser showing the strongest and more significant effect (**Figure 2B**). It has been proposed that MAMPs-perception elevates plant immunity against *Pst*DC3000 via compromising the onset of bacterial virulence (Anderson et al., 2014; Lovelace et al., 2018; Nobori et al., 2018). To assess the impact of AA on bacterial growth *in planta*, we tested the effect of Gln, Ser, Val and His on bacterial virulence gene expression after infiltration. To this end, we co-infiltrated *Pst*DC3000 with Gln, Ser, and Val individually in naïve plants and assessed the expression of the T3SS regulator *HrpL*, the T3E *AvrPto*, and the COR synthesis gene *cfl*. These three bacterial genes are induced at early stages of the infection and have been used as markers of virulence (Lovelace et al., 2018; Nobori et al., 2018). While Gln and Ser significantly suppressed the expression of *HrpL, AvrPto*, and *cfl*, Val only suppressed *HrpL, AvrPto* (**Figure 2C**). The co-infiltration of *Pst*DC3000 with His had no effect on the expression of these genes (**Figure 2D**), suggesting that increased growth does not depend on increased virulence. Overall, these data suggest that Gln and Ser may play an important role in suppressing bacterial virulence gene expression in flg22-treated plants. As a way of comparison, we tested the effect of citric acid, aspartic acid, and 4-benzoic acid on bacterial growth at 50 μM and 500 μM concentrations as previously reported by Anderson et al (Anderson et al., 2014) in seedlings. Our data revealed that the 500 μM combination of the three metabolites modestly increased bacterial growth in mock-treated leaves but had no effect on bacterial growth in flg22-treated leaves (**Figure S6A**). In addition, these metabolites did not affect the expression of T3SS in mock- or flg22-treated plants, suggesting that, unlike in seedlings, these metabolites do not induce virulence and bacterial growth *in planta* (**Figure S6B**).

**Figure 2.**
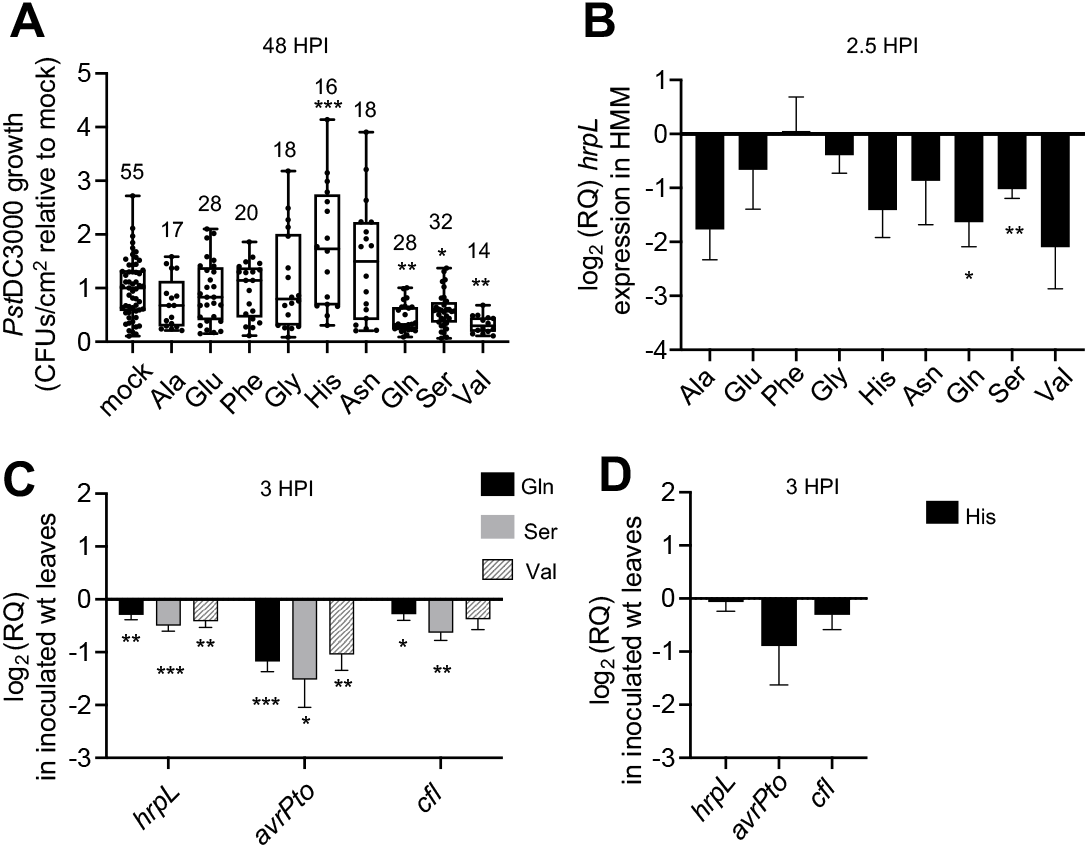
Amino acids that accumulate in MTI-elicited leaves modulate *Pst*DC3000 growth and virulence in naïve wild-type plants. (**A**) Bacterial growth 48 HPI of *Pst*DC3000 alone (mock) or co-infiltrated with individual amino acids in leaves of naïve plants and normalized by the growth of *Pst*DC3000 in mock. Numbers atop boxes indicate number of plants used per condition. (**B**) *hrpL* gene expression 2.5 h after inoculation of liquid HMM medium with *Pst*DC3000 supplemented with the indicated amino acids and normalized by the expression in non-supplemented HMM medium. (**C**) Expression of virulence genes in leaves of naïve plants 3 HPI with *Pst*DC3000 co-infiltrated with either Gln (n=12), Ser (n=12), or Val (n=18), normalized by the expression in non-supplemented *Pst*DC3000. (**D**) Expression of virulence genes in *Pst*DC3000 co-infiltrated with His in naïve plants 3 HPI (n=4). Statistic analysis: Brown-Forsythe and Welch ANOVA tests compared with mock (A); p values were adjusted by two-stage step-up false discovery rate method (Q = 0.01). One sample t-tests for (B)(C)(D). Combined data of 3 (A)(C)(D) and 4 (B) independent experiments.

Although effective at suppressing bacterial growth and virulence gene expression, the increase in AA concentrations is only one of the many responses elicited after flg22 perception in Arabidopsis. To compare the virulence-suppressing activity of Gln+Ser with that of flg22 pre-treated plants, we co-infiltrated *Pst*DC3000 with Gln+Ser in naïve wild-type plants or infiltrated *Pst*DC3000 alone in flg22 pre-treated plants, and assessed the expression of the *hrpL, AvrPto*, and *cfl*. While the expression of virulence genes was significantly lower in *Pst*DC3000 co-infiltrated with Gln+Ser compared to *Pst*DC3000 infiltrated alone in naïve plants (**Figure 3A**), they were expressed to similarly low levels in flg22-treated plants with or without Gln+Ser co-infiltration (**Figure 3B**). Importantly, both naïve plants and flg22 pre-treated plants co-infiltration with Gln+Ser suppressed the expression of virulence genes to a similar extent, suggesting that MTI suppresses bacterial virulence by means of the virulence-suppressing activity of Gln and Ser.

**Figure 3:**
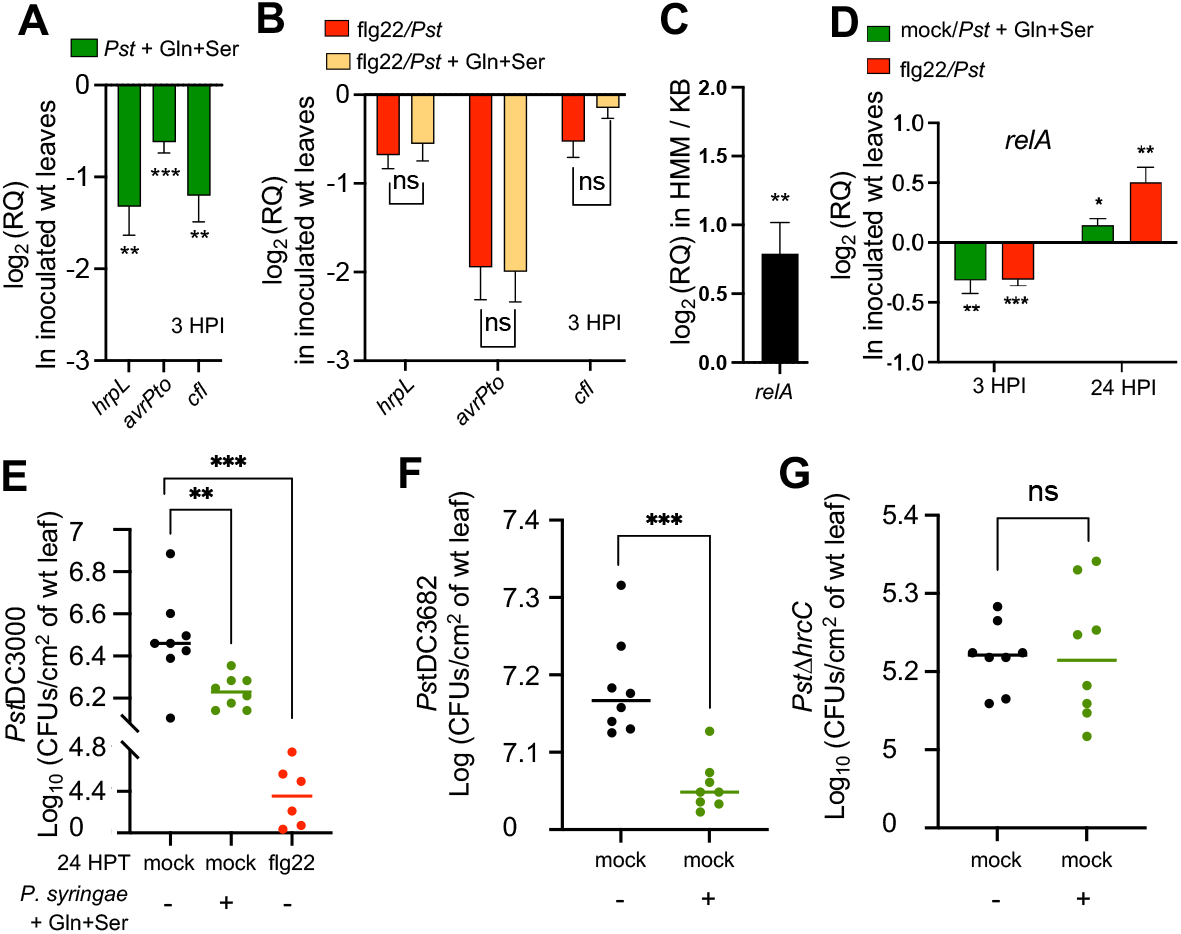
MTI suppresses bacterial virulence and growth via Gln and Ser accumulation. **(A)** Expression of virulence genes in naïve plants 3 HPI with *Pst*DC3000 and Gln+Ser (Mean *±* SEM, n=12). (**B**) Expression of virulence genes in flg22-treated plants 3 HPI with *Pst*DC3000 alone (n=36) or with *Pst*DC3000 and Gln+Ser (n=28); (Mean *±* SEM). (**C**) relA expression in hrp inducing minimal medium (HMM) normalized by its expression in King’s B rich medium (KB). (**D**) *relA e*xpression 3 HPI in mock-treated plants (green) with *Pst*DC3000 and Gln+Ser (n=27) or in flg22-treated (red) plants with *Pst*DC3000 alone (n=11)(Mean *±* SEM). (**E**) Bacterial growth 48 HPI in mock- (black) or flg22-treated (red) plants infiltrated with *Pst*DC3000 alone, or mock-treated plants (green) infiltrated with *Pst*DC3000 and Gln+Ser. (**F**) *Pst*DC3682 growth 48 HPI in naïve plants co-infiltrated with Gln+Ser (green). (**G**) *Pst*Δ*hrcC* growth 48 HPI in naïve plants infiltrated alone (black) or co-infiltrated with Gln+Ser (green). Statistic analysis: one-sample t-tests (A)(B)(C); Welch t-test (D); t-test (E)(F). (A)(B)(C) combination of 3 independent experiments. (D)(E)(F) one representative experiment of at lease 3 independent experiments.

To explore potential mechanistic connections between AA availability and *Pst*DC3000 virulence suppression, we tested the expression of *relA*, a bacterial gene that controls the onset of survival and virulence programs in response to nutrient starvation (Traxler et al., 2008). *relA* encodes a synthase/hydrolyze of the signaling metabolite guanosine penta- and tetra-phosphate, collectively known as (p)ppGpp. Like in other bacterial species, *Pst*DC3000 expressed *relA* in a nitrogen-poor minimal medium to higher levels than in a rich medium (**Figure 3C**). When co-infiltrated with Gln+Ser, *Pst*DC3000 expressed lower levels of *relA* than when infiltrated alone in naïve plants 3 HPI (**Figure 3D**). Similarly, *Pst*DC3000 expressed lower levels of *relA* in naïve plants than in flg22-treated plants 3 HPI. At 24 HPI, however, *Pst*DC3000 expressed higher levels of *relA* when co-infiltrated with Gln+Ser in naïve plants or infiltrated alone in flg22-treated plants (**Figure 3D)**. These data suggest that the extra availability of Gln and Ser early in the leaf colonization process keeps relA expression and bacterial (p)ppGpp concentrations below control levels, likely preventing the early onset of bacterial virulence, while causing excessive virulence expression at later time points when bacterial metabolism should be channeled towards producing growth instead of virulence.

To assess the contribution of Gln+Ser to MTI-mediated suppression of bacterial growth, we tested the growth of *Pst*DC3000 co-infiltrated with Gln+Ser in naïve plants side by side with *Pst*DC3000 inoculated alone in flg22-treated plants. Plants pre-treated with flg22 for 24 h prior to bacterial inoculation suppressed bacterial growth to a larger extent than Gln+Ser co-infiltration in naïve plants, suggesting that beyond Gln and Ser accumulation, other responses elicited by flg22 contribute to suppressing bacterial growth (**Figure 3E**). Like *Pst*DC3000, the growth of the COR synthesis mutant strain *Pst*DC3682 was also suppressed when co-infiltrated with Gln+Ser (**Figure 3F**). Unlike *Pst*DC3000, however, the growth of the T3SS mutant strain Δ*hrcC* was similarly low in both AA-supplemented and non-supplemented leaves (**Figure 3G**), suggesting that Gln+Ser suppresses bacterial growth primarily via suppressing the expression of the T3SS. In addition, the co-infiltration of *Pst*DC3000 with Gln+Ser in leaves of naïve *fls2* plants bypassed the need for flg22 perception to suppress *hrpL* and *cfl* expression (**Figure S7A)** and bacterial growth (**Figure S7B**). Overall, our data show that Gln+Ser supplementation does not elicit plant defense responses and that the expression of bacterial virulence is significantly lower when *Pst*DC3000 is co-infiltrated with Gln+Ser *in planta*, suggesting that increased AA concentrations contribute to bacterial growth-suppression directly by suppressing virulence rather than indirectly, by inducing plant defense responses.

### SA signaling contributes to the accumulation of amino acids in MAMP-elicited leaves

The Arabidopsis *SA-induced Deficient-2* (*sid2*) mutant is a loss-of-function mutant of *ICS1* that does not induce the synthesis of SA in response to microbial attack, a phenotype associated with attenuated responses to flg22 perception and enhanced susceptibility to *Pst*DC3000 infections (Wildermuth et al., 2001; Tsuda et al., 2008). SA signaling has been shown to positively regulate the expression of AA importers and the overall AA uptake activity in flg22-treated seedlings (Zhang et al., 2022). To infer if SA plays a similar role in the accumulation of AA in plants, we assessed the expression of AA/H^+^ symporters in flg22-infiltrated leaves of wild-type and *sid2-2* plants 8 HPT. Like in seedlings, while leaves of wild-type plants responded to flg22 perception by inducing several AA transporters, *sid2-2* only partially induced these genes (**Figure S9**), suggesting that SA plays an important signaling role in AA transport in mature leaves. Since SA is an important mediator of MAMP-elicited signaling and AA transport, we hypothesized that SA may also play an important role in the local accumulation of AA in MAMP-treated leaves. To test this hypothesis, we assessed the concentration of total AA in whole leaves and the leaf apoplast of wild-type and *sid2-2* plants 24 HPT with flg22. While leaf AA concentrations remained the same 24 HPT with water or flg22 (**Figure 4A**), an increase in AA concentration lower than in the wild type was detected in the leaf apoplast of flg22-treated *sid2-2* plants (**Figure 4B**), suggesting that SA-mediated signaling is required for leaves to accumulate AA in response to flg22 perception. To test for potential connections between MAMP-perception, SA-mediated responses, and bacterial virulence suppression, we tested if flg22-treated *sid2-2* plants were able to suppress *Pst*DC3000 virulence to a similar extent as wild-type plants. While flg22-pretreated wild-type plants significantly suppressed *hrpL* expression, *sid2-2* plants allowed *Pst*DC3000 to express *hrpL* to a similar extent as mock-treated *sid2-2* plants 1 HPI (**Figure 4C)**. By 3 HPI, however, flg22-pretreated *sid2-2* plants suppressed the expression of *hrpL* to the same extent as flg22-pretreated wilt-type plants (**Figure 4C**), showing a delayed suppression of virulence in *sid2-2* compared to wild type. To address the contribution of increased AA levels on bacterial virulence and growth independently of SA-mediated defense, we sought to bypass SA signaling by co-infiltrating *Pst*DC3000 with Gln+Ser in *sid2-2* plants. Like in wild-type or *fls2* plants 24 HPT with flg22, *Pst*DC3000 co-infiltrated with Gln+Ser expressed similarly low levels of *hrpL* and *cfl* in naïve *sid2-2* and naïve wild-type 3 HPI (**Figure 4D)**. In addition, the co-infiltration of *Pst*DC3000 with Gln+Ser significantly suppressed bacterial growth in naïve *sid2-2* plants (**Figure 4E**) albeit to a lower extent than in wild-type or *fls2* plants (**Figure S7**). These data suggest that SA-mediated signaling positively regulates the accumulation of AA in flg22-treated leaves, thus suppressing bacterial virulence at early time points after inoculation. The supplemented Gln+Ser, however, did not fully suppress *Pst*DC3000 growth in *sid2-2*, suggesting that in addition to suppressing the early onset of virulence, other defense mechanisms regulated by SA contribute to suppressing bacterial growth. Alternatively, the exogenous supplementation of Gln+Ser may not fully mimic the sustained availability of AA in the leaf apoplast MTI-elicited plants.

**Figure 4:**
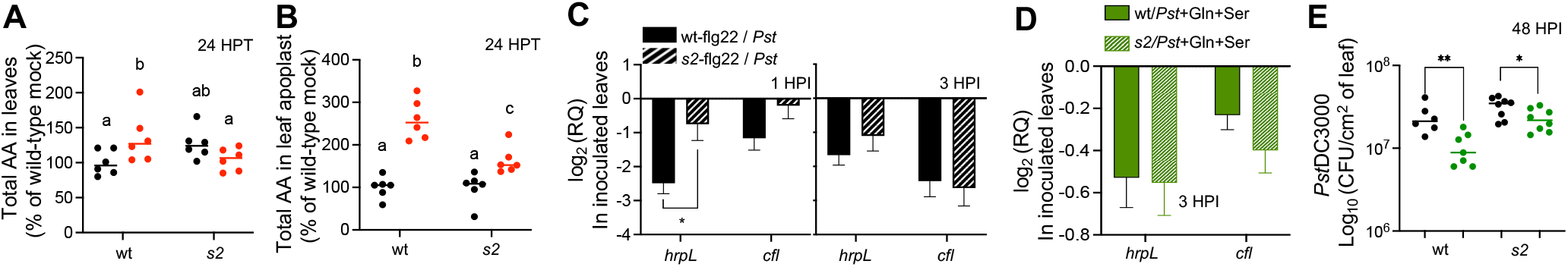
SA-mediated signaling contributes to flg22-induced accumulation of AA and bacterial virulence suppression. (**A**) Total AA in mock (black) or flg22-treated (red) leaves of wild-type (wt) or *sid2-2* (*s2*) plants 24 HPT. (**B**) Total AA in leaf apoplast of mock (black) or flg22-treated (red) wt or *s2* naïve plants 24 HPT. (**C**) Expression of virulence genes in wild-type or *s2* flg22-treated plants inoculated with *Pst*DC3000 alone and normalized by gene expression in mock-treated plants 1 HPI and 3 HPI; Mean *±* SEM (n=3); (**D**) Expression of virulence genes in wt or *s2* naïve plants co-infiltrated with *Pst*DC3000 and Gln+Ser; Mean *±* SEM (n=12). (**E**) Bacterial growth in wt or *s2* naïve plants 48 HPI with *Pst*DC3000 alone (black) or *Pst*DC3000 and Gln+Ser (green). Statistical analysis: one-way ANOVA (A)(B); t-test (C)(D)(E). (A)(B)(C)(E) one representative experiment from at lease 3 independent experiments. (D) combination of 3 independent experiments.

### High endogenous levels of amino acids partially bypass SA-mediated defense

To assess the expression of virulence genes and bacterial growth in conditions that better mimic the high concentrations of AA that *Pst*DC3000 encounters in the leaf apoplast of MTI-elicited wild-type plants, we sought to use naïve plants that accumulate high concentrations of AA constitutively without expressing the defense responses elicited by MAMPs. The Leucine-Histidine Transporter-1 (LHT1) transporter localizes at the plasma membrane of leaf mesophyll cells of mature leaves, where it takes up AA from the leaf apoplast (Hirner et al., 2006). *lht1-1* plants constitutively accumulate higher concentrations of AA in the apoplast of mature leaves, making this mutant a good candidate to test our hypothesis. However, *lht1-1* plants develop spontaneous cell death lesions, accumulate high concentrations of SA, and develop an enhanced disease-resistance phenotype against bacterial infections as they reach an age of 4 weeks (Liu et al., 2010). The pernicious effect of cell death and SA accumulation were fully rescued by the *sid2* mutation in the *sid2-2 x lht1-1* (*s2l1*) double mutant (Liu et al., 2010). However, the *sid2-2* mutation also eliminated the constitutive AA accumulation in leaf apoplast (**Figure S8**) making this genetic background unsuitable for testing our hypothesis. As an alternative to *lht1-1*, we tested AA levels in *gdu1*-1D, a gain-of-function mutant of the AA-export regulatory protein GLUTAMINE DUMPER-1 that accumulates high concentrations of AA in mature leaves (Pilot et al., 2004). Unlike *s2l1*, the double mutant *sid2-2* x *gdu1*-1D (*s2g1*), accumulates high concentrations of AA in mature leaves and leaf apoplast. While the apoplastic concentration of AA in *sid2-2* was similar to that of wild-type plants, it was higher in *s2g1* than in *sid2-2* or the wild type (**Figure 5A**). The expression of *hrpL, AvrPto*, and *cfl* was lower in *s2g1* compared to *sid2-2* (**Figure 5B**), reminiscent of the lower expression of these genes in naïve wild-type plants when *Pst*DC3000 was co-infiltrated with Gln+Ser compared to *Pst*DC3000 infiltrated alone (**Figure 3A**). In addition, and consistently with previous results obtained with Gln+Ser supplementation, bacterial growth was significantly lower in the *s2g1* double mutant compared to *sid2-2*, however not as low as when Gln+Ser was co-infiltrated with *Pst*DC3000 in wild-type plants (**Figure 5C**). Altogether, these data demonstrate that high Gln concentrations contribute to suppressing *Pst*DC3000 virulence and that additional SA-mediated defense mechanisms contribute to bacterial growth suppression.

**Figure 5.**
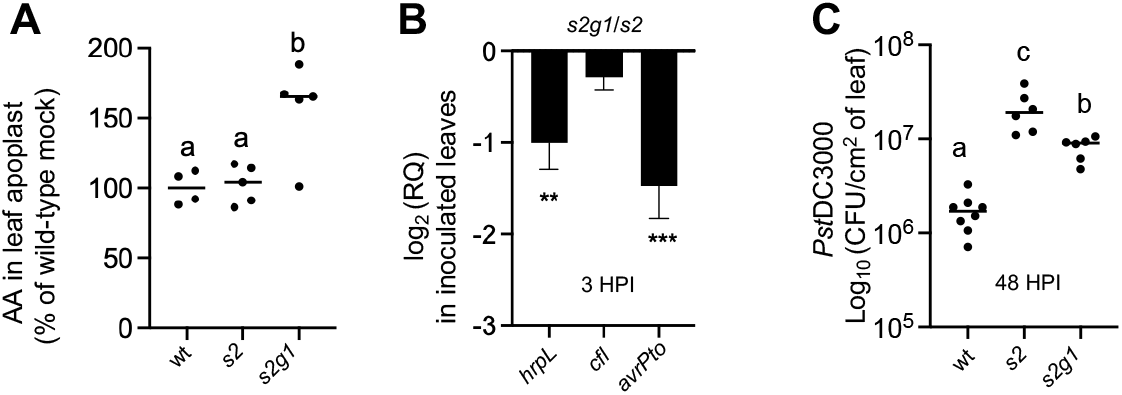
Increased endogenous levels of Gln impact *Pst*DC3000 virulence and growth in naïve plants. (**A**) Gln concentration in the leaf apoplast of wild-type wt, *s2*, and *s2g1* naïve plants. (**B**) Expression of virulence genes in *s2g1* naïve plants inoculated with *Pst*DC3000 alone and normalized by the expression in *s2* plants; Mean *±* SEM (*s2g1* n=15; *s2* n=13). (**C**) *Pst*DC3000 growth in wt, *s2*, and *s2g1* naïve plants 48 HPI. Statistical analysis: Brown-Forsythe and Welch ANOVA tests (A) and (C); p values were adjusted by the two-stage step-up false discovery rate method (Q = 0.01). (A) and (C) one representative experiment from at least three independent experiments. (B) combination of 3 independent experiments.

## Discussion

### Increased availability of amino acids in leaf apoplast suppresses the onset of bacterial survival responses and virulence

Plant bacterial pathogens must sense the environment and activate virulence pathways to suppress plant defense before they can colonize the leaf apoplast (Xin et al., 2018). While high levels of virulence may compromise growth by diverting energy away from the central metabolism, the expression of low levels of virulence may hinder bacteria’s ability to suppress plant defense responses (Xin et al., 2018). In most bacterial species, survival and growth are coordinated at least in part by the intracellular concentrations of (p)ppGpp (Traxler et al., 2008). In nitrogen-limiting, or otherwise suboptimal growth conditions (e.g., oxidative stress and osmotic stress) such as those experienced in minimal medium, bacteria express high levels of the (p)ppGpp-synthesis genes *relA*, thus allowing (p)ppGpp to accumulate. In turn, (p)ppGpp binds to RNA polymerase, shifting transcription towards genes that are essential for survival. In the leaf apoplast, the availability of AA and other sources of carbon and nitrogen changes as the infection progresses, demanding *Pst*DC3000 to constantly adjust (p)ppGpp levels to maximize growth. Upon entering the leaf apoplast, *Pst*DC3000 spends the first 3-6 hours sensing the new environment and adjusting its metabolism to express virulence genes in preparation for the rapid growth that takes place later in the infection (Nobori et al., 2018; Xin et al., 2018). In the early stages of host colonization, the main function of the T3SS is to suppress plant defense responses that would otherwise terminate the infection. Indeed, *Pst*DC3000 mutants compromised to express a functional T3SS, elicit strong MTI and fail to infect otherwise susceptible host plants (Hauck et al., 2003). Similarly, *Pst*DC3000 mutant strains compromised for the synthesis of (p)ppGpp, are unable to express wild-type levels of T3E proteins and only produce modest infections in otherwise susceptible Arabidopsis plants (Chatnaparat et al., 2015a; Chatnaparat et al., 2015b). The early expression of the T3SS has been proposed as an important factor to predict the success of *Pst*DC3000 to produce infections (Nobori et al., 2018). In agreement with previous studies, our data show that 3 HPI in MTI-elicited plants, *Pst*DC3000 expresses levels of *relA* and virulence genes lower than those detected in naïve plants. The low expression of *relA* in MTI-elicited plants is consistent with the metabolic adjustment to the nitrogen-rich leaf apoplast environment provided by plants that have accumulated AA prior to the inoculation (**Figure 1, Table I, Figure 3B, C**). Similarly, *Pst*DC3000 expresses low levels of *relA* and virulence genes when co-infiltrated with Gln+Ser into naïve wild-type plants (**Figure 2, Figure 3A, C**), again suggesting that high Gln+Ser concentrations, such as those encountered when infiltrated in leaves of MTI-elicited plants, serve a passive defensive function aimed at lowering bacterial stress responses and virulence. Twenty-four hours post-inoculation, however, *Pst*DC3000 co-infiltrated with Gln+Ser in naïve plants or infiltrated alone in MTI-elicited plants expressed higher levels of *relA* compared to bacteria infiltrated alone in naïve plants (**Figure 3C**). These data suggest that *Pst*DC3000 experiences a higher level of stress at later time points when co-infiltrated with Gln+Ser in naïve plants, perhaps due to not having been able to express virulence at higher levels to suppress plant defense responses earlier in the infection (**Figure 3A, 3C**). Importantly, *Pst*DC3000 expressed similarly low levels of T3SS, T3E, and COR biosynthesis genes when infiltrated into flg22-treated plants (**Figure 3A**) or co-infiltrated with Gln+Ser into naïve plants (**Figure 3B**), suggesting that Gln+Ser and flg22 pre-treatment suppress bacterial virulence gene expression through similar mechanisms and pathways.

### Amino acid-mediated suppression of virulence only partially suppresses bacterial growth

MTI suppresses not only the growth of the virulent *Pst*DC3000 wild-type strain but also that of T3SS mutants that typically produce mild infection in wild-type naïve Arabidopsis plants, suggesting that other mechanisms beyond T3SS suppression may account for the MTI-mediated protection against bacterial infections (Nobori et al., 2018). This evidence is consistent with the suppressing activity that AA exert on *Pst*DC3000 growth described in this study. AA supplementation in naïve plants did not suppress *Pst*DC3000 growth to the same extent as MTI-elicited plants did (**Figure 3E**). In addition, although effective at suppressing the growth of bacterial strains that do not produce coronatine (**Figure 3F**), AA supplementation did not suppress the growth of the Δ*hrcC* strain (**Figure 3G**). These data suggest that the increased availability of amino acids elicited by flg22 in wild-type plants suffices to suppress bacteria virulence but it is insufficient to fully suppress bacterial growth. This model is consistent with AA playing a defensive role in suppressing virulence, a critical mechanism in the early stages of an infection, but implies that other offensive responses are needed to further restrict bacterial growth in MTI elicited plants. This hypothesis is supported by the fact that, while AA suppressed bacterial virulence similarly in wild-type and *sid2-2* plants (**Figure 4D**), they only partially suppressed bacterial growth in *sid2-2* compared to wild-type plants (**Figure 4E**). In addition, while the expression of virulence genes is suppressed to a greater extent in *s2g1* than *sid2-2* (**Figure 5B**), the enhanced-disease susceptibility phenotype of *sid2-2* was only partially rescued by the increased availability of AA in the *s2g1* background (**Figure 5D)**. These data demonstrate that beyond the early suppression of virulence, active defense mechanisms mediated by SA-signaling contribute to restricting bacterial growth as the infection progresses. Hence, we propose that the bacterial infection level 48 HPI depends on both defensive responses that suppress virulence within the first 3 HPI and offensive responses that create a harsher environment as bacteria continue to colonize the leaf apoplast.

### SA-signaling positively regulates the accumulation of amino acids in response to MAMPs perception

SA regulates defense responses against biotrophic and hemibiotrophic pathogens at multiple levels. In addition to playing an essential role in the activation of cell-death mediated resistance initiated by plant intracellular receptors that detect bacterial T3E, SA plays a critical role in the activations of MTI upon MAMP perception (Tsuda et al., 2008). In a previous study, we unveiled that SA plays a positive role in regulating the basal update and the enhanced uptake of AA elicited by MAMPs-perception in Arabidopsis seedlings (Zhang et al., 2022). Like in seedlings, SA is necessary to induce the expression of AA/H^+^ symporters in response to MAMP-perception in mature leaves (**Figure S9**) (Zhang et al., 2022). The biological function of such induced gene expression and enhanced AA uptake activity detected in seedlings is still unclear. One possibility is that the enhanced uptake lowers the concentrations of AA in the leaf apoplast and deprives *Pst*DC3000 of carbon and nitrogen. In support of this possibility, the expression of *relA*, a bacterial proxy for nitrogen scarcity, was higher in *Pst*DC3000 infiltrated into MTI-elicited plants than in mock-treated plants 24 HPI (**Figure 3C**). In addition, the upregulation of *relA* 24 HPI will likely induce the onset of survival programs, diverting energy away from central metabolism and perhaps further compromising bacterial growth.

Since *sid2-2* plants accumulate less AA than wild-type plants in the leaf apoplast in response to flg22 treatment (Figure 4B) and take up less AA in response to flg22 perception in seedlings (Zhang et al., 2022), we hypothesize that the SA-dependent uptake of AA contributes to the overall accumulation of AA in flg22-treated leaves of wild-type plants. However, we cannot rule out the possibility that the higher concentrations of AA that build up in flg22-treated leaves originate from the up-regulation of cellular processes that contribute AA to the cellular pool or from the down-regulation of functions that consume AA. This hypothesis is in line with the increased concentration of AA in both leaf apoplast and phloem sap of flg22-treated leaves (**Figure 1D, E**), showing that AA accumulation results from increasing the intracellular pool of AA rather than the sequestration via phloem-upload shutdown. The transcriptional analysis presented in **Figure S2** suggests that flg22 induces proteolysis and senescence, two processes that contribute AA to the cellular pool. This transcriptional signature in mature leaves is consistent with previous studies showing that flg22 perception induces senescence in Arabidopsis seedlings (Denoux et al., 2008). In addition, flg22 perception suppresses processes that consume AA (e. g. aminoacyl tRNA Biosynthesis). Importantly, SA plays a positive role in regulating senescence and autophagy, suggesting that the compromised senescence and autophagy of *sid2-2* plants could explain the compromised accumulation of AA in response to flg22 perception (**Figure 4A,B**).

### How amino acids suppress bacterial virulence during MTI

A model of bacterial virulence suppression is presented in Figure S10. When plants perceive MAMPs and mount defense responses in the absence of bacterial interference, the series of events that follow MAMP perception may well represent the sequence of events that follow the perception of non-virulent bacterial strains: MAMP perception elicits changes in the metabolite composition of the leaf apoplast that prevent bacterial proliferation. Data obtained in a previous study showed the perception of flg22 or elf26, shuts down the uptake of AA within the first 4 HPT in Arabidopsis seedlings (Zhang et al., 2022). We hypothesized that this inhibition of AA uptake could be the direct result of the alkalinization of the leaf apoplast as shown in the liquid medium of liquid-grown seedlings (Gómez-Gómez et al., 1999). Due to technical limitations, a similar increase in the pH of leaf apoplast of plants has so far not been demonstrated. In addition, technical limitations hinder the assessment of AA uptake in the leaf apoplast of mature leaves directly. Thus, we can only infer that flg22 perception inhibits the uptake activity of AA/H^+^ symporters via the alkalinization of the leaf apoplast, which would lead to the accumulation of AA outside the mesophyll cells. We have detected an increase in AA concentrations in the leaf apoplast of flg22-treated leaves at 24 HPT but not earlier (**Figure 1C**). This increased concentration of extracellular AA concurs with the accumulation of AA over a 24 h period after flg22 treatment (**Figure 1B**). In seedlings, the early inhibition of uptake elicited by MAMPs subsides between 4 HPT and 8 HPT and is replaced by an enhanced AA uptake activity that lasts several hours (Zhang et al., 2022). This increased uptake of AA is mediated by the AA/H^+^ symporters LHT1. The perception of flg22 induces the expression of *LHT1* and other AA/H^+^ symporters in an SA-dependent manner in both seedlings and mature leaves (**Figure S9**). However, the induced expression of the AA transporters does not prevent AA concentrations from increasing in the leaf apoplast of mature leaves, suggesting that any increase in import activity might be offset by an increased export of AA into the leaf apoplast. In seedlings, flg22 perception inhibited the uptake of AA and concurrently induced the expression of AA/H^+^ symporters well ahead of the increased AA uptake activity, suggesting that the inhibition of AA uptake offsets any increase in the number of transporters that localize to the plasma membrane. We do not know if flg22 perception also inhibits the uptake of AA in mature leaves or how long the inhibition lasts. However, the prolonged inhibition of uptake alongside an increased availability of intracellular AA that constantly leak out of the cells via AA facilitators would likely produce the accumulation of AA detected 24 HPT with flg22. Further studies will be necessary to understand what metabolic processes yield intracellular AA that leak out of the cells and into the apoplast, as well as the identity of transporter proteins that coordinate their activity to move AA out of the plant cells.

## Materials and Methods

### Plant growth conditions

Arabidopsis thaliana plants were grown on pellets (Jiffy Pellets #7) in growth Conviron Gen100 chambers with 9 hours of light photoperiod, 100 μE.m^-2^.s^-1^ light energy, 23ºC, and 70% humidity. Plants were watered with half-strength Hoagland solution three times a week for the first four weeks. Then, plants were watered with tap water three times a week for two more weeks. Wild-type plants were the reference Col-0 ecotype unless otherwise specified. The *sid2-2* mutant was a gift from Mary Wildermuth at the University of California Berkeley. Seedlings were grown in liquid Murashige and Skoog (MS) basal medium with vitamins (Phytotechnology Laboratories) supplemented with 0.5 g/L MES hydrate and 0.5% sucrose, pH 5.7 corrected with KOH, in a Conviron Adaptis A1000 growth chambers (Conviron, Inc.) under 16 h of light photoperiod, 23ºC constant temperature, 100 μE.m^-2^.s^-1^ light energy, and 80% relative humidity.

### Extraction and quantification of amino acids with fluorometric assay

The fourth and fifth pairs of leaves were collected in 2 mL round-bottom microcentrifuge tubes and flash-frozen at the indicated time points (see figure legends) after infiltration. For seedlings, 1 μM flg22 was added on day 10 and 24 hours later samples were flash-frozen in liquid nitrogen. Frozen leaf and seedling samples were then lyophilized and weighed. The dry weight was adjusted to approximately 11 mg in each sample and crushed into powder with two 5 mm stainless steel beads with Qiagen TissueLyser for 5 minutes at 3000 strokes per minute. Extraction in 400 μL of 80% methanol was performed by mixing thoroughly with Qiagen TissueLyser for 5 minutes at 3000 strokes per minute. The samples were incubated at 28ºC overnight, centrifuged at 17.000 x g for 10 minutes, and 300 μL of the supernatant were transferred to a new tube and mixed with 200 μL of chloroform and 100 μL of water. The samples were mixed with a Qiagen TissueLyser for 1 min at 3000 strokes per minute and then centrifuged at 17.000 x g for 10 minutes. The clear upper phase (150 μL) was transferred to a new tube for air drying and resuspended in HPLC grade water.

Apoplastic washing fluids were obtained using the infiltration and centrifugation method (Lohaus et al 2001). The fourth and fifth pairs of leaves were infiltrated with water or 1 μM flg22 as described above. Twenty-four hours post infiltration, leaves from two plants (eight leaves total) were pooled together as one sample, rolled together to fit inside the cylinder of a 5 mL sterile syringe. The tip of the syringe was sealed with parafilm, the leaves inside the cylinder were submerged in sterile water, the plunger was inserted in the cylinder and the leaves were infiltrated by repeatedly applying positive and negative pressure by the moving the plunger up and down. Once fully infiltrated, the leaves were removed from the syringe and pat dry on tissue paper to remove surface water. Leaves of 1 sample were stacked again together and put inside the cylinder of a clean 5 mL syringe without a plunger. The syringe was inserted in a clean 15 mL conical tube and centrifuge for 5 min at 500 x g in a swinging bucket clinical centrifuge. Apoplastic washing fluids were collected at the bottom of the 15 mL conical tube, transferred to 1.5 mL microcentrifuge tubes and stored at -80 ºC. Phloem sap was collected as described by Corbersier et. al. 2002 (REF) with minor modification. Briefly, previously infiltrated leaves were detached from the plant, the petioles were rinsed in 10 mM EDTA to remove cellular content at the edge of the cut, and submerged in 400 μL of 5 mM EDTA at 100% RH for 6 h in the dark at 23ºC. Samples were then lyophilized and stored at room temp. L-Amino acids were quantified with the commercial “L-AA fluorometric assay kit” (Bio Vision, Inc) following the manufacturer’s directions.

### Profiling of amino acids

Fourth and fifth pairs of leaves were infiltrated with water or 1 μM flg22 for 24 hours before being collected in 2 mL round bottom tubes and flash-frozen. Samples were then lyophilized and weighed. The dry weight of each sample was adjusted to approximately 5 mg. Dried leaves were crushed into powder with metal beads in a Qiagen TissueLyser. Amino acids were extracted with 200 μL of HCl 10 mM spiked with 0.1 mM Novarline (Sigma-Aldrich) as an internal extraction control, plus 200 μl of Chloroform. Each sample was vortexed for 2 minutes and centrifuged at 17,000 x g for 5 minutes. The supernatant was transferred to a new tube for analysis using a method previously described (Collakova et al). Briefly, samples were derivatized with AccQ•Tag™ labeling fluorescence kit (Waters Corporation) and amino acids were separated using Ultra-Performance Liquid-Chromatography (UPLC) with an Acquity UPLC and detected using an Acquity-UPLC FLP fluorescence detector (Waters Corporation).

### Bacterial virulence expression

*Pst*DC3000 from an overnight liquid King’s B culture was transferred to a sterile 50 mL conical tube with fresh media (1:10 dilution) and incubated at 28ºC for 2 hours with agitation. For *in vitro* gene expression analysis, *Pst*DC3000 was pelleted by centrifugation, washed three times with sterile water, and resuspended in Hrp-inducing minimal medium (Huynh et al, 1989). Bacteria were added to wells of a 12-well tissue culture plate at a final optical density (OD_600nm_) of 0.2 and individual amino acids were added at a 5 mM final concentration to the corresponding wells by triplicate. The plates were incubated at room temperature (23ºC) for 150-190 minutes before the bacteria were transferred to microcentrifuge tubes, pelleted, and stored at -80ºC for RNA extraction. For *in planta* gene expression analysis, *Pst*DC3000 was resuspended at an OD_600nm_ of 0.2 in 5 mM MES (pH 5.7) for control samples. For control experiments, *Pst*DC3000 was infiltrated into leaves as is. For amino acids supplementation experiments, amino acids were added to the bacterial suspension final concentration and immediately infiltrated into the fourth and fifth pairs of leaves. Leaf samples were taken between 150 to 180 minutes after infiltration, flash-frozen in liquid nitrogen, and stored at -80ºC for RNA extraction.

### *In planta* bacterial infection assays

*In planta* infections were performed as described previously by Hauck et al (A Pseudomonas syringae type III effector suppresses cell wall-based extracellular defense in susceptible Arabidopsis plants) with minor modifications. Plants were grown in pit pellets and watered as indicated above. Fully expanded leaves (fourth and fifth pairs) of six-weeks-old plants were pressure infiltrated from the abaxial side of the leaf lamina with a sterile 1 mL needleless syringe loaded with bacterial suspension. *Pst*DC3000 inoculum from -80ºC glycerol stock was streaked on a LB-agar plate and incubated overnight at 28ºC. A small amount of bacterial lawn from the overnight plate was transferred to 3 mL King’s Broth media in a 50 mL conical tube and incubated under agitation for 3-4 hours at 28ºC to an OD_600nm_ of 0.4 to 0.7. Bacteria were pelleted by centrifugation at 8.000 x g for 3 min and resuspended in sterile water. The pelleting and resuspension process was repeated three times to wash the bacterial inoculum. Bacteria were then resuspended in 5 mM MES buffer (pH 5.7) and the OD_600nm_ was adjusted to 0.0002. For *Pst*Δ*hrpC* (T3SS mutant) and *Pst*DC3682 (coronatine mutant), an OD_600nm_ of 0.02 was used for plant inoculation to compensate for the slow *Pst*Δ*hrpC* and *Pst*DC3682 bacterial growth in wild-type Col-0 plants. For amino acids supplementation, amino acids were added to the inoculum suspension to reach 10 mM final concentration. For the elicitation of MTI prior to bacterial inoculation, leaves were infiltrated with water (mock) or with 1 μM flg22 peptide via pressure infiltration (as described above) 24 hours prior to bacterial inoculation. Infections were allowed to proceed for 48 h. For bacterial infection quantification, leaves were removed from the plants and 5 mm in diameter leaf disc were collected with a hole puncher and transferred to 2 mL round bottom tubes. Samples were ground in 400 μL of sterile water with metal beads in a Qiagen TissueLyser. Ten-fold serial dilutions of the ground leaf samples were plated on LB rifampicin 50 μg/mL Omin Tray™ single-well rectangular plates (Nunc™) and incubated overnight at 28ºC. Colony forming units (CFU) were counted under the microscope (Olympus SZ61) and informed as CFUs per cm^2^ of leaf area.

### Plant gene expression analysis

Fourth and fifth pairs of leaves from 6-week-old plants were pre-treated with water or 1 μM flg22 as indicated in the text. 10 mM Gln and Ser were buffered in 5 mM MES and infiltrated alone or with *Pst*DC3000 at an OD_600nm_ of 0.2. Samples were taken between 150 to 180 minutes after infiltration of leaves and snap frozen in liquid nitrogen and stored at -80ºC for RNA extraction.

### RNA extraction and RT-qPCR

Leaf samples or bacterial pellets were removed from the -80ºC freezer and kept on dry ice until grinding. Right before grinding, Trizol Reagent (Invitrogen) was added to each tube and samples were ground with metal beads and a Tissue Lyser (QIAGEN). For bacteria RNA extraction either from leaves or bacterial pellets, Trizol Reagent and 0.1 mm in diameter Zirconia/Silica beads (Sigma-Aldrich) were added to each tube and pulverized in a bead beater (BioSpec Products) for 2 min at 7.000 stroke/min before proceeding with RNA extraction. RNA samples were then treated with DNAse I (Promega) to remove genomic DNA contamination. cDNA was synthesized with M-MLV retro-transcriptase (Promega) and random hexamers primers (Invitrogen). qPCR reactions were carried out with SYBR Green reagent (CoWin Biosciences, Inc.) in an Applied Biosystems™ 7500 Fast Real-Time PCR Instrument. *Actin2* (At3g18780) expression and *gyrA* (PSPTO_1745) were used as normalizing housekeeping genes for *Arabidopsis* and *Pst*DC3000, respectively. Primer sequences were: gyrA-F: 5’-ttcaatgctgatcccggaagaagg-3’; gyrA-R: 5’-atttcctcaccatccagcacctga-3’; hrpL-F: 5’-tcaggaaagctgggaagacgaagt-3’; hrpL-R: 5’-atgttcgacggcaggcaatcaatg-3’; avrPto-F: 5’-atgacgggagcgtcaggaatcaat-3’; avrPto-R: 5’-atccgttcgggttcatagtcgcaa-3’; cfl-F: 5’-tgctcgtctcgtcgccaa-3’; cfl-R: 5’-cgatacccttagttagtcctgtgg-3’; CYP81F2-F: 5’-ctcatgctcagtatgatgc-3’; CYP81F2-R: 5’-ctccaatcttctcgtctatc-3’; MYB51-F: 5’-ctctcttcacgcccttcacg-3’; MYB51-R: 5’-ccggaggttatgcccttgtg-3’; ICS1-F: 5’-gaactcaaatctcaacctcc-3’; ICS1-R: 5’-actgcgacgagagaagaaac-3’; PR1-F: 5’-ttcttccctcgaaagctcaa-3’; PR1-R: 5’-aaggcccaccagagtgtatg-3’

### *In silico* analysis of Affymetrix gene expression data

Gene sets within GO terms pertaining to amino acid metabolism were obtained from TAIR (The Arabidopsis Information Resource). Shorter lists of genes were made by clustering genes based on functions that contribute (amino acids biosynthesis, autophagy, senescence, and proteolysis) or consume (Glucosinolates biosynthesis, Phenylpropanoids biosynthesis, Aminoacyl tRNA biosynthesis) free amino acids. Gene expression values for each gene were retrieved from the TAIR Affymetrix DataSet 1008080727 (Brunner and Nürberger 2005; NCBI-Gene Expression Omnibus GSE561). This dataset consist of three replicates of water (mock) and flg22-infiltrated leaf samples taken 1 HPT and 4 HPT. Induced (>1.5-fold change) or suppressed (<1.5-fold change) genes (p 0.05 t-test) belonging to the sources (contribute) or sinks (consume) of amino acids were used to estimate their overall contribution to the pool of free amino acids. The average gene expression of each cluster was calculated to build the heat map shown in Figure S2.

### Quantification of radiolabeled AA in leaves and leaf apoplast

Using a needle-less 1 mL syringe, the 2^nd^ and 3^rd^ pair of leaves from six-week-old plants were infiltrated with sterile water or 1 μM flg22. Twenty-four hours later, the same leaves were infiltrated with a mix of 1 mM cold amino acids (Gln, Ser, or Pro) spiked with the corresponding ^14^C labeled amino acid. One hour after infiltration, the leaves were removed from the plants at the base of the lamina, quickly rinse in water, and transferred to a 5 mL sterile syringe without a plunger. Twelve leaves were rolled together and inserted with the petiole upwards into the syringe. The exit of the syringe was sealed with parafilm, the syringe was filled up with 0.24 M Sorbitol, the plunger was inserted, and the leaves were infiltrated following cycles of positive and negative pressure by moving the plunger up and down until complete infiltration. The leaves were removed from the syringe, surface dried with a paper towel, and transferred to a clean 5 mL syringe without a plunger with the base of the petiole facing upwards. The syringe with the leaves was transferred to a 15 mL conical tube, centrifuged at 500 X g for 3 min, and AWF was collected at the bottom of the 15 mL conical tube. The twelve leaves in each sample were transferred to a 2 mL round bottom ultracentrifuge tube containing two 5 mm metal beads and 400 μL of 5 % bleach (to remove chlorophyll) and ground with the TissueLyser. The recovered AWF, typically 100 μL, was mixed with 100 μL of scintillation cocktail and kept in the dark for 2 h. One hundred microliters of the leave’s lysates were lyophilized overnight, resuspended in 100 μL of 0.24M Sorbitol, mixed with 100 μL of scintillation cocktail, and stored in the dark for 2 h. Radioactivity in each sample was estimated as counts per minute (CPMs) using a Wallac 1450 TriLux MicroBeta 96-well plate scintillation counter (PerkinElmer).

---

**Figure S1.**
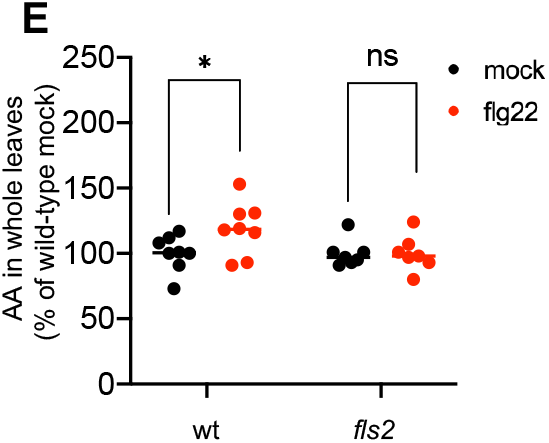
Total amino acids in whole leaves of wt and *fls2* plants infiltrated with water (black) or flg22 (red) 24 HPT.

**Figure S2.**
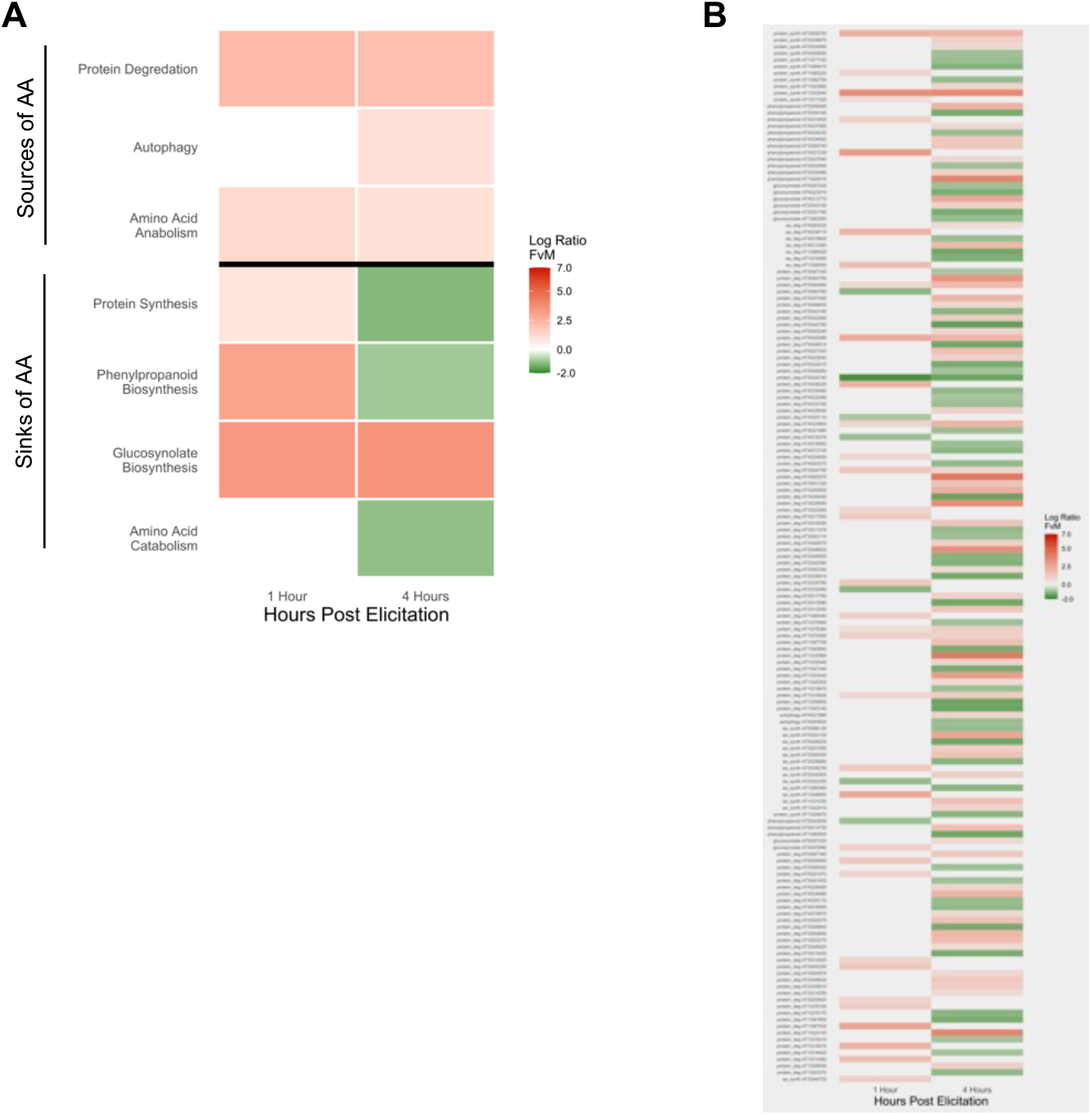
Expression of flg22-responsive Arabidopsis genes with roles in amino acids metabolism. (A) Average gene expression1 HPT and 4 HPT with 1 μM flg22 in \leaves of wild-type Col-0 plants normalized by mock treatment. Genes were organized by processes that contribute or consume AA. (B) Expression of individual genes induced (>1.5 fold) or suppressed (<1.5 fold) in response to flg22 perception 1 and 4 HPT. Only genes whose expression passed the t-test p 0.05 value were included. Affymetrix data was obtained from Brunner and Nürberger 2005; TAIR ExpressionSet 1008080727.

**Figure S3:**
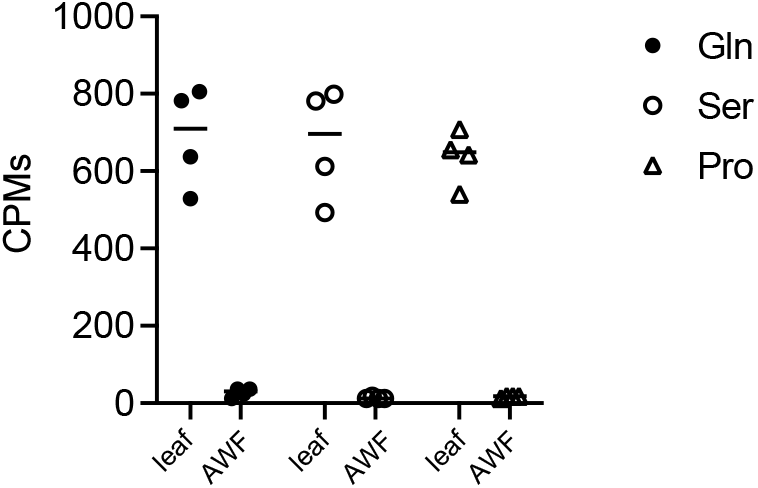
Externally supplied amino acids were quickly taken up by plant cells. Leaves were infiltrated with ^14C^Gln (squares), ^14C^Ser (red), or ^14C^Pro (triangles), each one supplemented with 1 mM of the corresponding cold amino acid. One hour later, the leaves were infiltrated again with 0.24 M sorbitol and centrifuges to obtain apoplastic washing fluids (AWF). After centrifugation, both whole leaves (leaf) and AWF were processed to assessed and assess counts per minute (CPMs) CPMs. Two independent experiments yield identical results.

**Figure S4:**
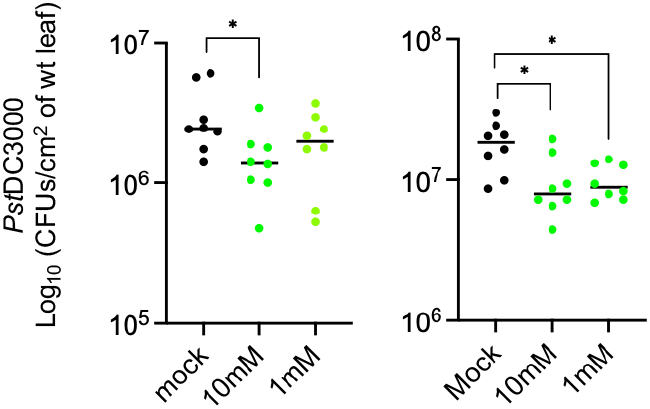
Gln + Ser suppress PstDC3000 growth. Bacteria were inoculated in naïve wild-type plants alone (black) or co-infiltrated with Gln+Ser at two different concentrations. CFUs were counted 48 HPI. Shown are two independent experiments. Data analysis: t-test.

**Figure S5:**
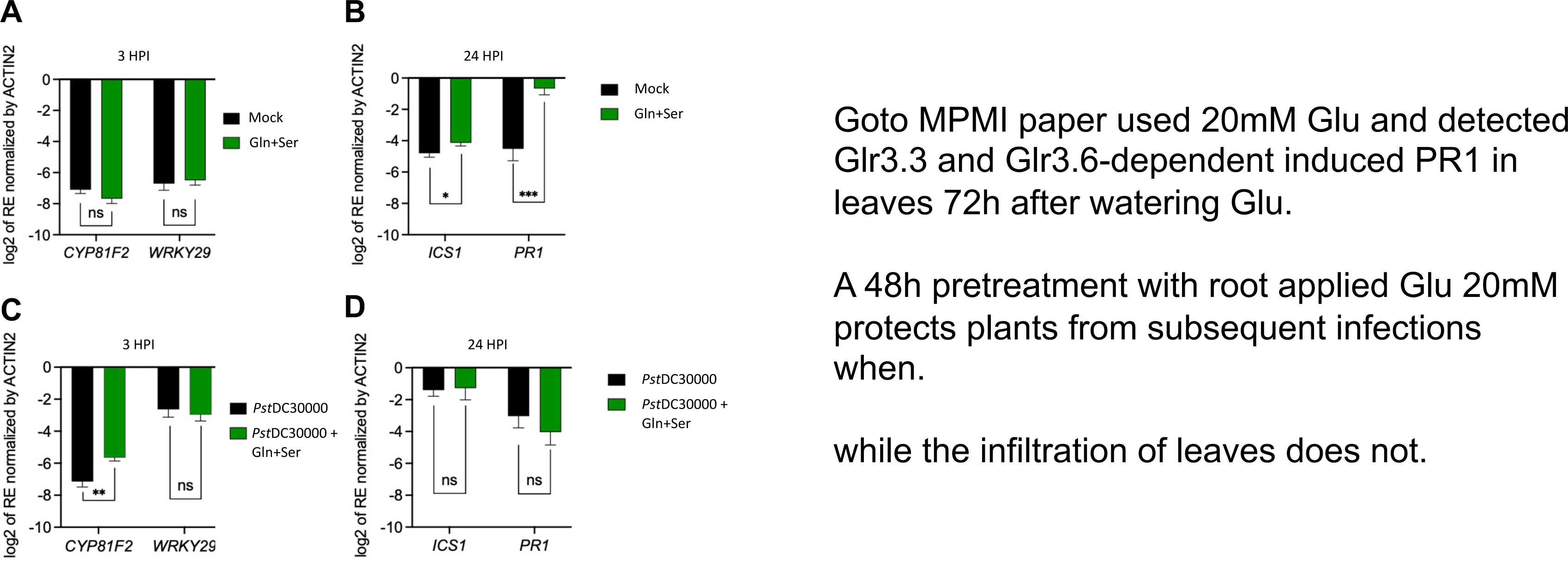
Arabidopsis defense gene expression in response to Gln+Ser infiltration. Arabidopsis gene expression 3 h (**A**) or 24 h (**B**) after infiltration of Gln+Ser, normalized by expression in water-infiltrated leaves. Arabidopsis gene expression 3 h (**C**) or 24 h (**D**) post-co-infiltration of *Pst*DC3000 with Gln+Ser normalized by the expression in *Pst*DC3000 infiltrated alone. Data analysis: Welch t-test. Results are a combination of 3 independent experiments with n=9 (A), n=12 (B), n=11 (C and D).

**Figure S6:**
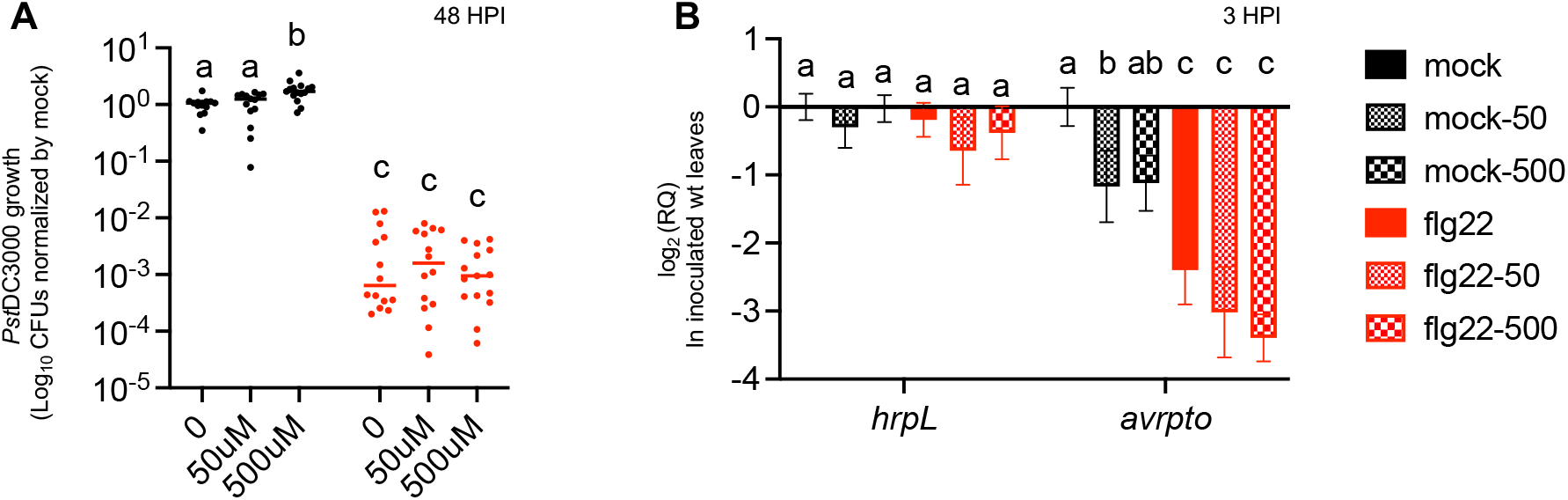
Effect of citric acid, aspartic acid, and 4-benzoic (CAB) on PstDC3000 growth and virulence in adult plants. (**A**) Bacterial growth in mock- (black) or flg22-treated (red) plants infiltrated alone (0) or co-infiltrated with either 50 μM or 500 μM of CAB. (**B**) Bacterial gene expression 3 HPI in mock- (black) or flg22-treated (red) wild-type plants inoculated with *Pst*DC3000 alone (mock) or co-infiltrated with 50 μM or 500 μM of CAB.

**Figure S7.**
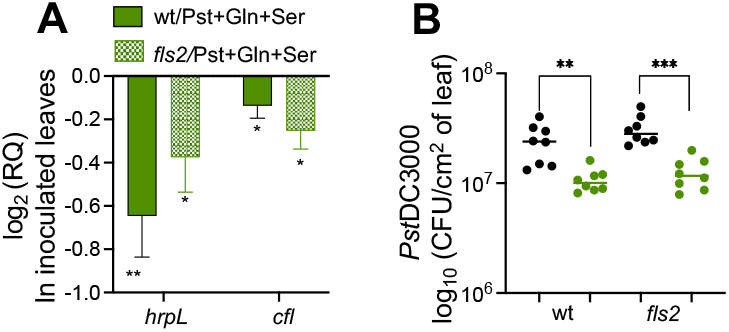
Exogenous supplementation of Gln+Ser bypasses flg22 perception to suppress bacterial virulence. (**A**) Bacterial gene expression 3 HPI in naïve wild-type (green solid) or *fls2* (green checker) plants co-infiltrated with *Pst*DC3000 and Gln+Ser. (**B**) Bacterial growth 48 HPI in naïve wild-type of *fls2* plants inoculated with *Pst*DC3000 alone (black) or co-infiltrated with Gln+Ser (green).

**Figure S8.**
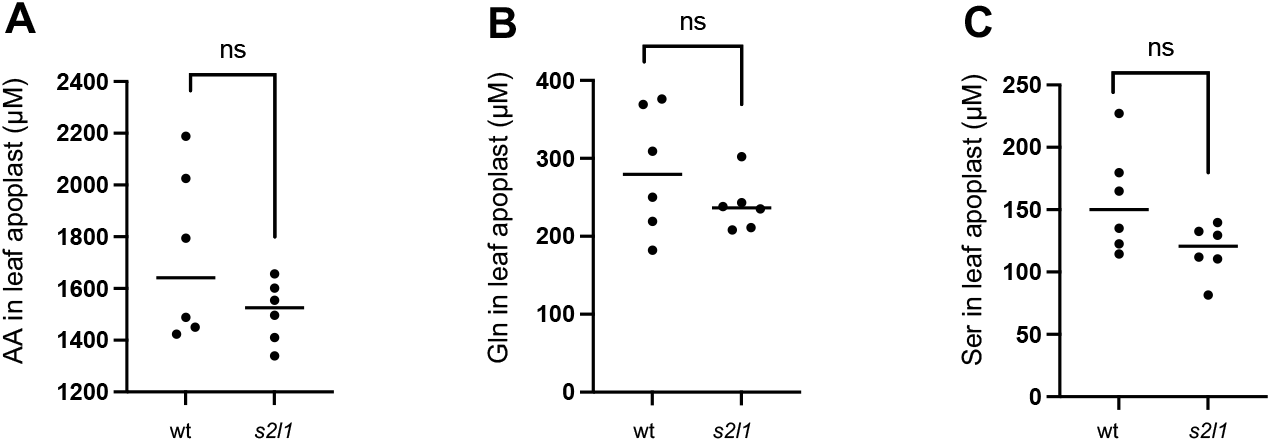
Salicylic acid is required for lht1 to accumulate AA in leaf apoplast. AA (A), Gln (B), Ser (C) concentrations in leaf apoplast of six-week-old wt and sid2-2 x lht1-1 (s2l1) plants.

**Figure S9:**
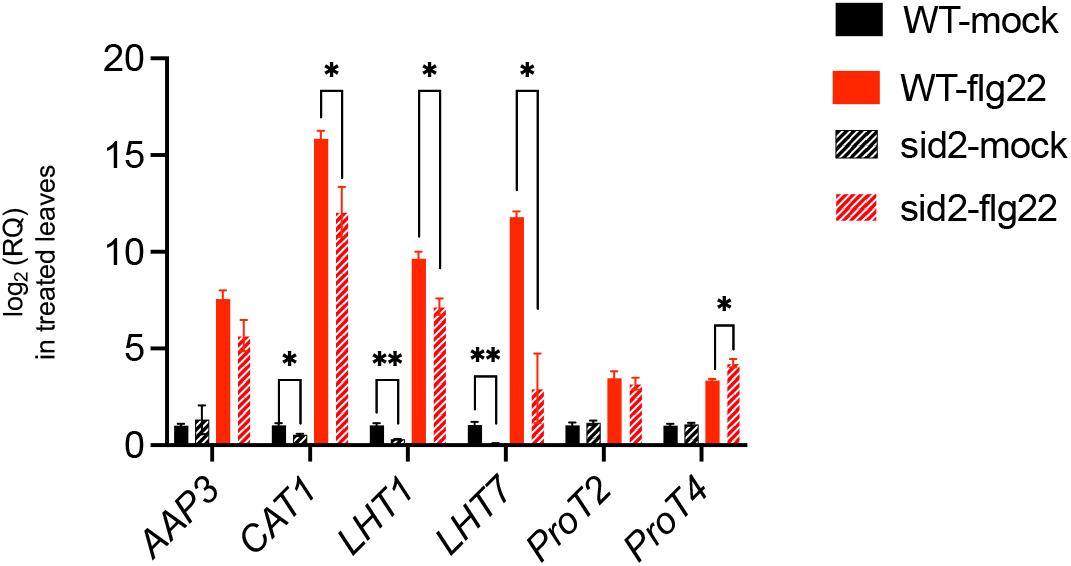
Amino acids transporters gene expression in flg22-infiltrated plants 8 HPT. Mean *±* SEM of wild-type (n=9) and *sid2-2* plants (n=9) RNA quantified by nanoString RNA hybridization analysis. Data analysis: t-test. Combination of 3 independent experiments.

**Figure S10:**
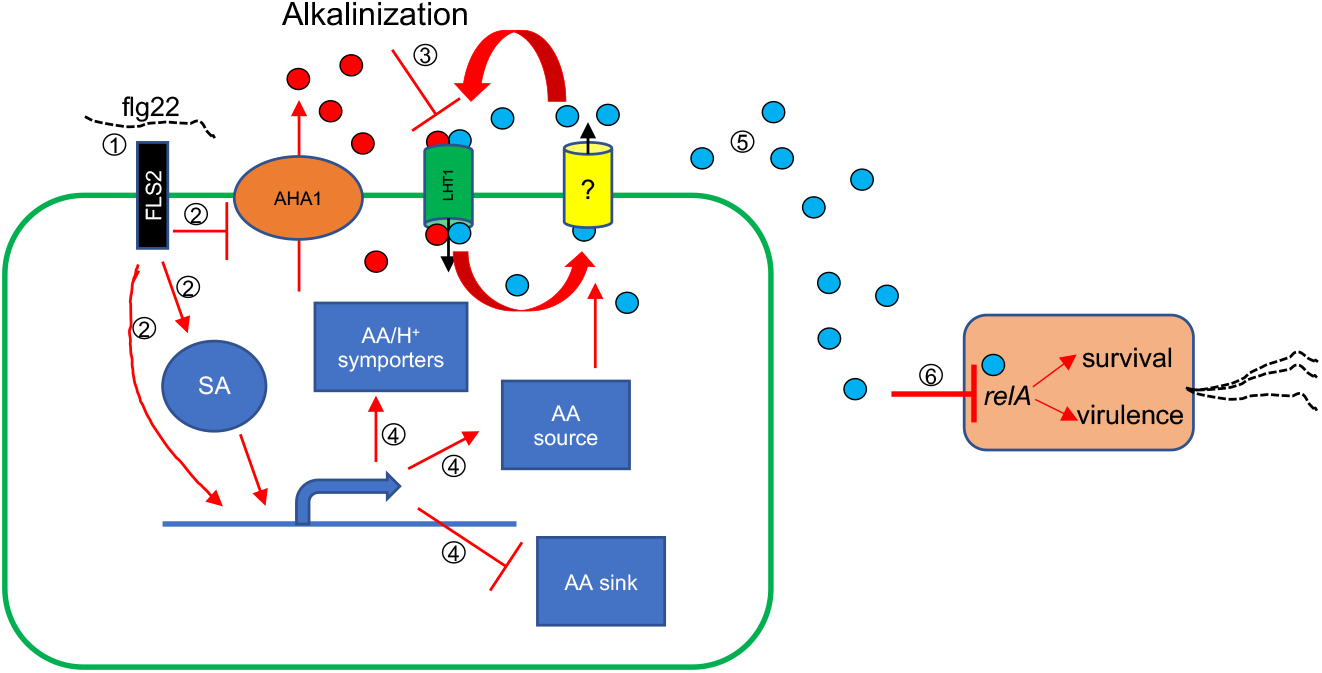
Proposed model: how amino acids suppress bacterial virulence during MTI. The perception of flg22 (1) activates of SA-dependent and independent signaling pathways induction (2) and suppresses the activity of the plasma membrane-localized H^+^-pump ATPase AHA1 (2), leading to the alkalinization of the apoplast and the inhibition of AA uptake (3). Meanwhile, signaling pathways activated by flg22 perception induce genes encoding AA/H+ symporters and others involved in cellular processes that enlarge the intracellular pool of AA (4) concomitantly with the suppression of genes involved in cellular processes that consume AA (4). Both the inhibition of uptake and the increased availability of intracellular AA, lead to the increased concentration of AA in the apoplast (5) and the suppression of *relA* (6), thus suppressing the onset of bacterial virulence and survival programs. H^+^ ions and AA are depicted as red and light blue circles, respectively.

